# A special role for anterior cingulate cortex, but not orbitofrontal cortex or basolateral amygdala, in choices involving information

**DOI:** 10.1101/2023.08.03.551514

**Authors:** Valeria V. González, Sonya A. Ashikyan, Yifan Zhang, Anne Rickard, Ibrahim Yassine, Juan Luis Romero-Sosa, Aaron P. Blaisdell, Alicia Izquierdo

## Abstract

Subjects often are willing to pay a cost for information. In a procedure that promotes paradoxical choices, animals choose between a richer option followed by a cue that is rewarded 50% of the time (No-info) *vs* a leaner option followed by one of two cues that signal certain outcomes: one always rewarded (100%), and the other never rewarded, 0% (Info). Since decisions involve comparing the subjective value of options after integrating all their features, preference for information may rely on cortico-amygdalar circuitry. To test this, male and female rats were prepared with bilateral inhibitory DREADDs in the anterior cingulate cortex (ACC), orbitofrontal cortex (OFC), basolateral amygdala (BLA), or null virus (control). We inhibited these regions after stable preference was acquired. We found that inhibition of ACC destabilized choice preference in female rats without affecting latency to choose or response rate to cues. A logistic regression fit revealed that the previous choice strongly predicted preference in control animals, but not in female rats following ACC inhibition. The results reveal a causal, sex-dependent role for ACC in decisions involving information.

---

The environment is full of unpredictable events, and information that reduces uncertainty about such events allows an organism to better predict and prepare for the future. However, several psychiatric conditions are characterized by a strong preference for information, or an Intolerance of Uncertainty, including Autism Spectrum Disorder, Substance Use Disorders, Attention-Deficit Hyperactivity Disorder, Generalized Anxiety Disorder, and Obsessive-Compulsive Disorder (Tolin, Abramowitz et al. 2003, Dugas, Buhr et al. 2004, Boulter, Freeston et al. 2014, Jenkinson, Milne et al. 2020, Mandali, Sethi et al. 2021). Intolerance of uncertainty has been linked to ‘pathological doubt’ during which the preference for information is dramatically increased (Tolin, Abramowitz et al. 2003).

Obtaining information can be crucial for survival. However, if information cannot be used to modify action, a bias toward information can be considered paradoxical or even suboptimal because organisms should not invest resources to obtain information that does not affect the outcome of a choice. Imagine a situation in which an organism chooses between two sources of delayed reward; if rewards are signaled before each choice, it will be worth investing in that information to choose the best option. In contrast, if the outcomes are signaled *after* the choice, information is useless, since the organism cannot change its choice. The latter has been broadly studied at the behavioral level. In the so-called paradoxical or suboptimal choice task (Stagner and Zentall 2010, McDevitt, Dunn et al. 2016, González, Izquierdo et al. 2023), animals are presented with two alternatives, one providing a lower rate of reinforcement with different stimuli indicating the presence (S+) or absence (S-) of delayed food (i.e. Info), and another one (S3) providing a higher rate of reinforcement but with non-differential stimuli signaling food (i.e. No- Info) (See **Figure 1A**). Birds consistently prefer the leaner but informative option despite the difference in reinforcement rates between alternatives (Stagner and Zentall 2010, Fortes, Vasconcelos et al. 2016, Macias, Gonzalez et al. 2021). However, rats more often show high variability, with some experiments showing preference for the No-Info option (Orduña 2015, Orduña 2016) and some showing preference for the Info option (Cunningham and Shahan 2019, Ajuwon, Ojeda et al. 2021). We propose that such individual differences allow us to test if animals use different strategies to make decisions, which can also shed light on how value is assigned.

**Figure 1.**
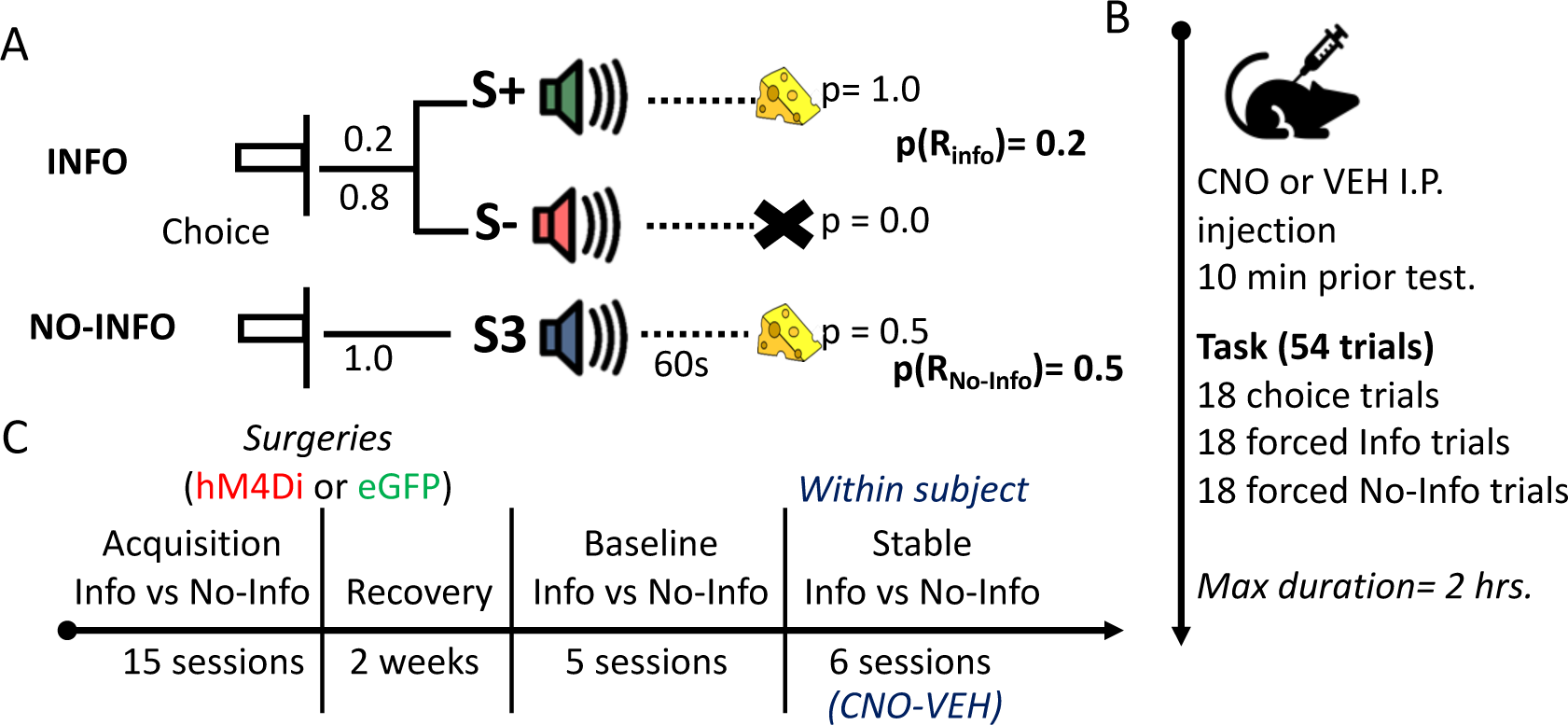
***Task structure and experimental timeline.* (A)** Rats choose between two levers (left or right). After pressing once, both levers retract and an auditory cue commences. If the rat chooses the ‘Info’ alternative, on 20% of the trials, tone S+ plays for 60s always ending with the delivery of one sugar pellet; the other 80% of the trials sound S- plays for 60s always ending without food. If the rat chooses the ‘No- Info’ option, a third sound S3 plays for 60s ending with the delivery of one sugar pellet in 50% of the trials. (**B**) CNO or VEH was administered 10 min before starting the task. Each session consisted of 54 trials in which rats received 18 choice trials (both levers available to choose), 18 forced Info trials (only the lever associated with the Informative alternative was available) and 18 forced No-Info trials (only the No-Info lever was available). Animals were required to complete the session in 2 hrs. (**C**) Rats were trained on the task over 15 sessions before they underwent bilateral viral infusion of inhibitory hM4Di DREADDs or null virus enhanced Green Fluorescent Protein (eGFP) in ACC, BLA, or OFC. After two weeks of recovery, baseline performance was reestablished for 5 sessions before administrating CNO and VEH (order counterbalanced) for three consecutive sessions with a wash-out day between drugs.

The neural substrates of reinforcement uncertainty have been broadly researched, with evidence pointing to a distributed network that involves prefrontal cortex (PFC) (Rushworth and Behrens 2008), striatum, hippocampus, basolateral amygdala (BLA) and mediodorsal thalamus (Winstanley and Floresco 2016, Soltani and Izquierdo 2019). Nevertheless, to our knowledge there is no investigation of the specific brain regions using this behaviorally well-documented paradoxical choice procedure.

Additionally, compared to primates (Iigaya, Story et al. 2016, Iigaya, Hauser et al. 2020), there is a paucity of rodent studies on the value of non-instrumental information and its neural substrates. When human subjects are assessed on choices between two cued alternatives- an informative vs. a non-informative one (but with no difference in overall reinforcement rate), subjects reliably prefer advanced information and Blood-oxygen-level-dependent (BOLD) signal in ventromedial PFC tracks the value of the anticipation of reward (Iigaya, Story et al. 2016, Iigaya, Hauser et al. 2020). Neural correlates of uncertainty have been found in different subregions of the PFC in several species, among them, orbitofrontal cortex (OFC) and anterior cingulate cortex (ACC) (Rushworth and Behrens 2008,

Bromberg-Martin and Hikosaka 2009, Wallis 2012, Blanchard, Hayden et al. 2015, Bromberg-Martin and Monosov 2020). Electrophysiological recording studies in rats and monkeys demonstrate that activity in OFC is associated with stimulus (cue) value and expected uncertainty, or probabilistic risk (Jo and Jung 2016, Riceberg and Shapiro 2017, Namboodiri, Otis et al. 2019, Jenni, Symonds et al. 2023). Similarly, studies have shown that ACC neurons signal value, uncertainty of predictions about rewards or punishments (Monosov 2017, Jezzini, Bromberg-Martin et al. 2021), and track trial-by-trial outcomes of choice (Procyk, Tanaka et al. 2000, Shidara and Richmond 2002). On the other hand, in probabilistic- discounting tasks, when rodents are required to select between a large uncertain option versus a small certain option, inactivation of BLA decreases the likelihood of choosing the uncertain option (Ghods-Sharifi, St Onge et al. 2009, Stopper and Floresco 2011, St Onge, Stopper et al. 2012). This suggests the contribution of BLA may be biasing choice towards larger rewards, especially when the delivery of these rewards is uncertain. Furthermore, selectively disrupting PFC-to-BLA connections increases choice of the larger yet increasingly uncertain reward, indicating that communication between these two regions serves to modify choice biases (St Onge, Stopper et al. 2012). Finally, lesions to either OFC or BLA (or their connection), results in slower learning about which option has the better payout, suggesting that these areas form a functional circuit for the adaptation of reward-maximizing strategies (Zeeb and Winstanley 2011, Zeeb and Winstanley 2013).

In this experiment, we examined the specific contributions of ACC, OFC, and BLA to this seemingly ‘suboptimal’ choice phenomenon via chemogenetic manipulation. On the additional evidence that these regions participate in decision confidence under uncertainty (Lak, Costa et al. 2014, Stolyarova, Rakhshan et al. 2019), we hypothesize that they may play vital, yet dissociable roles in decisions involving information value. To study this, we inactivated these regions during stable preference. We found that inhibition of ACC, but not OFC or BLA, destabilized choices involving information in females and reduced their ability to use previous choices to guide current decisions.

## Materials and Methods

### Animals

Sixty-six Long Evans rats (*Rattus Norvegicus*), 36 females and 30 males, acquired from Envigo served as subjects. Subjects were between aged post-natal days (PND) 90 and 140 at the start of the experiment. Subjects were pair-housed before and single-housed after surgeries in transparent plastic tubs with wood shaving bedding in a vivarium maintained on a reverse 12-hr light cycle. Experiments were conducted during the dark portion of the cycle at a minimum of five days per week. A progressive food restriction schedule was imposed prior to the beginning of the experiment to maintain rats at 85% of their initial free-feeding weights. Water was always available in their home-cages. The procedures used in this experiment were conducted under approval and following the guidelines established by the Chancellor’s Animal Research Committee at UCLA.

### Viral Constructs

To express DREADDs on putative projection neurons in ACC, OFC, or BLA, an adeno-associated virus AAV8 driving the hM4Di-mCherry sequence under the CaMKIIa promoter was used (AAV8-CaMKIIa-hM4D(Gi)-mCherry, packaged by Addgene, viral prep #50477-AAV8), thus targeting pyramidal neurons. A virus without the hM4Di DREADD gene but containing the fluorescent tag enhanced Green Fluorescent Protein, eGFP (AAV8-CaMKIIa-EGFP, packaged by Addgene, viral prep #50469-AAV8) was infused into ACC, OFC, or BLA as a null virus control. This null virus allowed us to control for non-specific effects of surgical procedures (i.e., craniotomy, anesthesia), exposure to AAV8, and non-specific effects of drug or injections.

### Behavioral apparatus

This experiment was conducted using operant testing chambers, measuring 30 x 25 x 20 cm (L x W x H). Each chamber was housed in separate sound- and light- attenuating environmental isolation chests (ENV-008, Med Associates, Georgia, VT). The front and back walls and ceiling of the chambers were constructed of clear Plexiglas, the side walls were made of aluminum, and the floors were built of stainless-steel rods measuring 0.5 cm in diameter, spaced 1.5 cm center-to-center.

Each chamber had a pellet-dispenser (ENV-203-45, Med Associates) and a cup type pellet receptacle (ENV-200R1M, Med Associates). When activated, one sucrose pellet was delivered into the cup. The opening of the cup was equipped with an infrared beam and photodetector to record entries into the food niche. A 3.5 cm wide operant lever was positioned one cm to the left and right of the food niche on the metal wall.

A speaker (ENV-224DM) on the ceiling of the chamber delivered a siren (cycling between 1500 and 1900 Hz at a 0.5 s rate), a 1000-Hz tone and a white noise, all were 8 dB above background to serve as the initial S+, S- and S3; and another siren (cycling between 4000 and 3500 Hz at a 0.5 s rate), a 3000- Hz tone and a click train (4/s) 8 dB above background served as a second set of S+, S- and S3 cues, counterbalanced across subjects. A diffuse incandescent light (ENV-227M, Med Associates) was located on the bottom panel of the right-side chamber wall, 6 cm from the ceiling.

### Surgical procedures

After completing the training phase, rats were anesthetized with isoflurane for bilateral infusion of ACC, OFC or BLA inhibitory (Gi) DREADDS (AAV8-CaMKIIα-hM4D(Gi)-mCherry, Addgene, Cambridge, MA, viral prep #50477-AAV8) or eGFP (AAV8-CaMKIIa-EGFP, Addgene, Cambridge, MA, viral prep #50469-AAV8). Craniotomies were created, and a 26-gauge guide cannula (PlasticsOne, Roanoke, VA) with a dummy injector was lowered, after which the dummy injector was replaced with a 33-gauge internal injector (PlasticsOne, Roanoke, VA) was inserted. Animals were infused with two bilateral sites of injections in BLA (0.2 µL at 0.1 µL/min in AP: -2.5, ML: ±5.0, DV: -8.6, and 0.1 µL at 0.1 µL/min in AP: -2.5, ML: ±5.0, DV: -8.3; total volume per side = 0.3 µL), two bilateral sites in OFC (0.15 µL at 0.1 µL/min in AP: 4.0, ML: ±2.5, DV: -4.4, and 0.2 µL at 0.1 µL/min in AP: 3.7, ML: ±2.5, DV: -4.6; total volume per side = 0.35 µL) and one bilateral site in ACC (0.3 µL at 0.1 µL/min AP: +3.7, ML: ±0.8, DV: -2.4 for a total volume of 0.3 µL per side). All measurements were taken from bregma.

After infusion, the cannula was left in place for ten additional minutes to allow diffusion.

### Drug treatment

Before testing, rats were given intraperitoneal (i.p.) injections of vehicle (VEH: 95% saline + 5% DMSO) or clozapine-N-oxide, CNO (3 mg/kg CNO in 95% saline + 5% DMSO) 10 min prior to beginning the behavioral task. The time was shorter than some other work (Hart, Blair et al. 2020, Ye, Romero-Sosa et al. 2023) to account for the longer duration of behavioral testing sessions as in a previous experiment (Stolyarova, Rakhshan et al. 2019). CNO and VEH were administered during *Stable Info vs No Info* condition in a within-subject design (**Figure 1B**), where animals received three sessions of CNO (or VEH) followed by a washout day with no injection or training, and then three sessions of VEH (or CNO) followed by another washout day, such that the order of drug administration was counterbalanced across animals.

### Behavioral procedure

#### Pretraining

Before training, ten sucrose pellets were given in the rat home cage to avoid food neophobia. On the first day, rats were trained to eat pellets from the pellet tray by delivering one pellet every 20 ± 15 s in the chamber (actual intertrial interval (ITI) values = 5, 10, 15, 20, 25, 30, and 35 s) for a total of 40 pellets. On days 2 and 3, rats were trained in an autoshaping procedure to lever press the left and right lever (lever presented in alternate order). Reinforcements were delivered following each lever press (i.e., a continuous reinforcement schedule) for a maximum of 40 pellets within a session, or after 30 min had elapsed.

#### Training of the Paradoxical choice task

The training stage of the task was comprised of two types of trials: choice and forced trials (**Figure 1C**). In choice trials, rats were required to choose between two levers available simultaneously. A choice trial started with the simultaneous insertion of both levers and the houselight turning on, the levers remained extended until a choice was made (no time limit). A choice was made by pressing a lever one time. After the choice was made, if the animal chose the Info alternative, both levers retracted and the houselight turned off. If the lever associated with the Info option was pressed, 20% of the time an auditory cue (S+) was presented for 60 s always ending with food; the other 80% of the time another auditory cue (S-) was on for 60 s never ending with food. The total percentage of reinforced trials was 20%. If the rat chose the No-Info alternative, a third auditory cue (S3) was presented for 60 s and ended with food on half of the trials. The percentage of reinforcement on this alternative was 50%. On forced trials, only one lever was presented at a time, following the same contingencies described above. All trials were separated by a 10-s intertrial-interval (ITI) during which all stimuli were off. A single session constituted 54 trials, 18 choice trials, and 36 forced trials. The maximum duration of a session was set to 120 min. The assignment of sounds to S+, S- or S3 and the lever side for each option was counterbalanced across animals but remained the same for individual animals throughout training. This condition lasted 15 sessions.

#### Stable preference

Approximately two weeks after surgery, rats received five training sessions as described above as a reminder of the task, used as a measure of baseline preference (Baseline Info vs No Info, **Figure 1B**). After this, rats received three sessions of the task with an i.p. injection of CNO and three sessions with an injection of VEH (order counterbalanced across animals). After the three-session round of injections, animals underwent a washout day in which they were not tested on the task. This condition lasted six sessions (eight days).

### Histology

At the conclusion of the experiment, rats were euthanized by an overdose of sodium pentobarbital (Euthasol, 0.8 mL, i.p.; VetOne, Paris, France) and transcardially perfused with phosphate buffered saline (PBS) followed by 10% buffered formalin acetate. Brains were extracted and post-fixed in this solution for 24 hours followed by 30% sucrose cryoprotection. Tissue was sectioned in 40-µM thick slices and cover slipped with DAPI mounting medium (Prolong gold, Invitrogen, Carlsbad, CA) visualized using a BZ-X710 microscope (Keyence, Itasca, IL), and analyzed with BZ-X Viewer software (**Figure 2A**).

**Figure 2.**
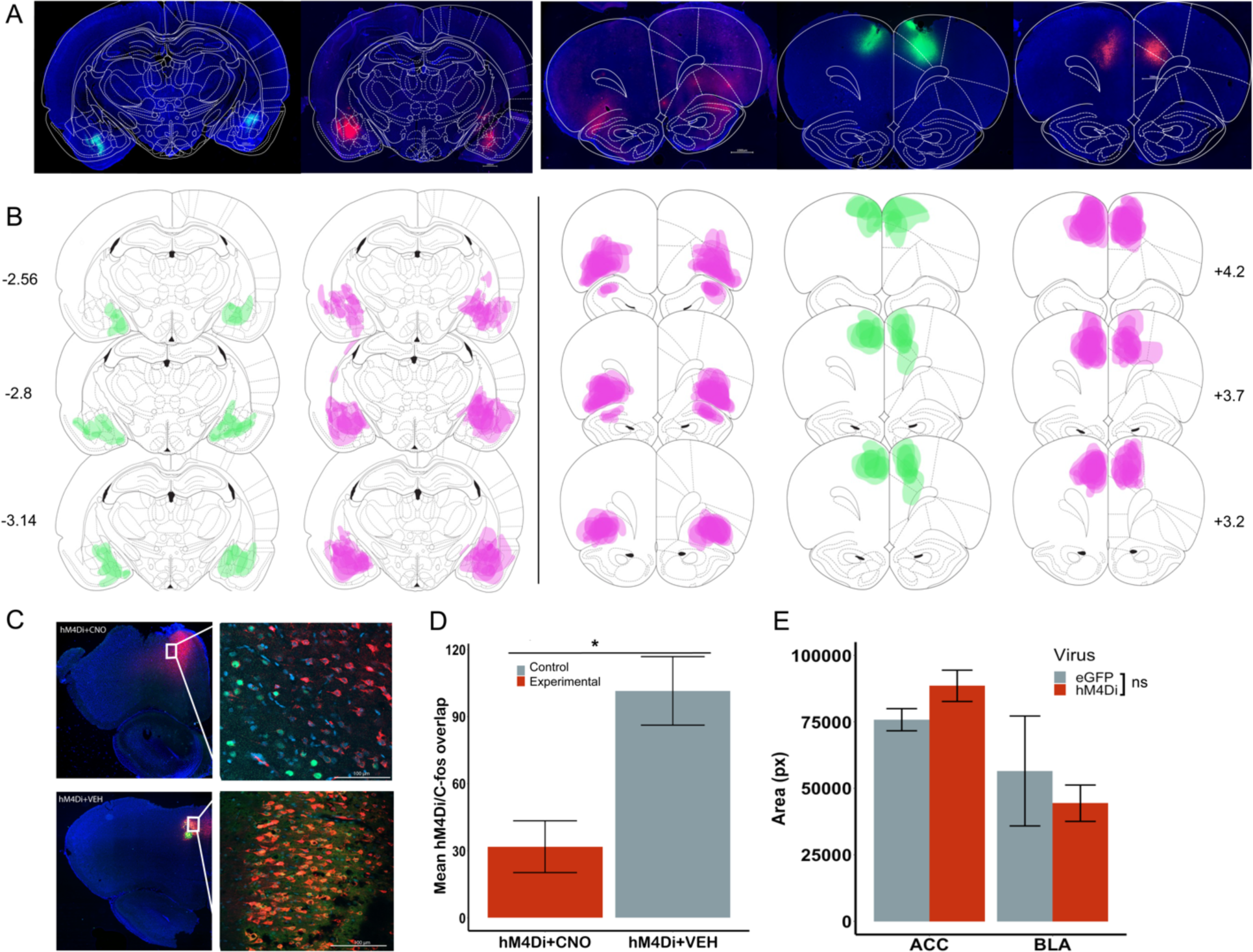
***Inhibitory DREADDs in ACC, BLA and OFC; Validation of ACC inhibition via c-fos immunohistochemistry.* (A)** Representative placement of inhibitory hM4Di DREADDs and eGFP null virus at Anterior-Posterior (AP) level +3.7 for ACC and OFC, and -2.8 for BLA relative to Bregma. Note, there were no eGFP for OFC. **(B)** Reconstructions of placement of inhibitory hM4Di DREADDs (pink) and eGFP null virus (*green*) at AP level +4.2, +3.7 and +3.2 for ACC, OFC, and AP -2.56, -2.8 and -3.14 for BLA relative to Bregma. **(C)** Representative image showing DAPI (blue), hM4Di-mCherry (red), c-fos immunoreactivity (green) and their overlap after injections of CNO and VEH in ACC at 2.5x and 20x. **(D)** Mean cell count of 4 images per condition, hM4Di+CNO, hM4Di+VEH for ACC. **(E)** Mean spread of viral expression across brain regions, pixel quantification was done using ImageJ software. ns= nonsignificant, *p<0.05, Mann-Whitney U test.

We have previously validated the efficacy of our CNO-activated inhibitory DREADDs *ex-vivo* in slice (Stolyarova, Rakhshan et al. 2019, Aguirre, Woo et al. 2023), *in vivo* using electrophysiological recordings (Ye et al. 2023), and behaviorally (Stolyarova, Rakhshan et al. 2019, Hart, Blair et al. 2020, Aguirre, Woo et al. 2023, Ye, Romero-Sosa et al. 2023) in OFC, ACC, and/or BLA. A group of ACC animals (N= 4) received a CNO or VEH injection 30 min prior to the beginning of the perfusion. In this group of animals, a subset of the 40 μm coronal sections were also stained for c-Fos, following an adapted Abcam protocol for dry-mounted slides (Schneider Gasser, Straub et al. 2006). In this protocol, mounted tissue was marked with a hydrophobic pen, any medium was added with a pippette and removed using a vacuum. The tissue was incubated for 22-24 h at 4 °C in a solution of primary antibody (1:2000 rabbit polyclonal to c-Fos, Abcam, Cambridge, MA) with 5% normal goat serum (Abcam, Cambridge, MA), and 0.1% TritonX-100 (Sigma, St. Louis, MO) in PBS. After the incubation, the tissue was washed with PBS 3 times during a 5-min period, then the brain slides incubated in a secondary antibody for 90 min protected from light at room temperature (0.1% TritonX-100, 5% normal goat serum, PBS solution with 1:500 goat anti-rabbit Alexa 488; Abcam, Cambridge, MA). Slides were washed again as described above. Then, tissue was incubated during 3 min with quenching reagent (1:1 ratio) to reduce background. Slides were washed with PBS and then cover-slipped with fluoroshield DAPI mounting medium (Abcam, Cambridge, MA).c-Fos inmunoreactivity quantification images were visualized with a 20x objective with a 724 μm by 543 μm field of view using a confocal microscope (Model LSM 900, Zeiss, Germany). For each region, four images were taken from two or three separate coronal sections from both hemispheres at the same approximate AP coordinate (ACC +3.7 mm). To verify DREADD-mediated inhibition of pyramidal-neurons in ACC after CNO or VEH administration we compared the number of c-fos–positive cells in the hM4Di-expressing regions to the number of c-fos–positive cells in neighboring (non-hM4Di- expressing, DAPI) areas. Cell counts were conducted using ImageJ software (**Figure 2C**). We found greater overlap number of c-fos positive cells in hM4Di+VEH than hM4Di+CNO cell areas (Mann-Whitney U test: W= 0, *p* = 0.009). We did not find a difference between the c-fos positive cells in DAPI areas (DAPI+CNO vs. DAPI+VEH; Mann Whitney U test: W= 8.5, p = 0.61) (**Figure 2D**). DREADDs and eGFP expression were determined by matching histological sections to a standard rat brain atlas (Paxinos and Watson, 2007) and quantifying fluorescence using ImageJ (Rueden, Schindelin et al. 2017) where two independent raters measured the area of fluorescent pixels for each animal per hemisphere (**Figure 2E)**.

To approximate the amount of viral expression observed in the tissue across brain regions (ACC, OFC and BLA) and virus (hM4Di and eGFP), we quantified the max area of fluorescent pixels of every animal presented in the reconstructions (**Figure 2B**). We used an independent samples t-test to compare viral spread (average fluorescent pixel area) for ACC hM4Di (MeanACC hM4Di= 88542.87) vs. ACC eGFP (MeanACC eGFP= 75795.5). The same analysis compared BLA hM4Di (MeanBLA hM4Di= 44399.54) and BLA eGFP (MeanBLA eGFP= 56519.33). No comparison with eGFP was conducted for OFC hM4Di (MeanOFC hM4Di= 100132) given the absence of eGFP animals in OFC. However, there were no significant differences between ACC and OFC hM4Di expression (p=0.26), and we show below that control groups were no different from each other and could be collapsed into one group. We found no significant differences in the viral expression between DREADDs hM4Di and control eGFP for ACC (*t*(18.52)= - 1.77, *p* = 0.093, -95% CI [27850, 2355]) or for BLA *t*(2.46)= 0.56, *p* = 0.624, 95% CI [-66729, 90969]).

### Data Analysis

All analyses were performed via custom-written code in MATLAB (MathWorks, Inc., Natick, MA). There were 3 main conditions in our analyses: 1) preference for the Info alternative (Info choice divided by the total number of choices) 2) latency to choose (the time from the beginning of the trial until a lever press was made), and 3) response rate (the number of entries into the food niche) during the 60-s cue duration.

Learning and performance data were analyzed with a series of mixed-effects General Linear Models (GLMs) (*fitglme* function; Statistics and Machine Learning Toolbox) first in omnibus analyses that included all factors (drug, virus and sex), with all groups (ACC, BLA, OFC and Control). Analyses were further pursued pending significant interactions in the full model. All post-hoc tests were corrected for the number of comparisons (with Bonferroni tests).

We also fit a logistic regression, a type of GLM for binary classification, to each rat’s data that belonged to either an experimental group or a control condition, separately by sex. We aimed to predict the probability of choice using 3 features: 1) Latency to choose, 2) Previous choice, and 3) Previous reward, depending on which option was chosen. This analysis was performed in a more fine-grained manner, on trial-by-trial data. We used only free-choice data for the outcome variable, and the previous choice and reward as either from free-choice or forced-choice data. The full model was *γ ∼ [1 + PrevChoice + PrevReward + PrevLatency]*, with a binomial distribution. A note that the β coefficients here correspond to log(odds ratio), which transforms odds ratio (>0) to a real number indicating how much one unit increase of the predictor contributes to log odds of the event occurring (versus not occurring). Values smaller than 1 correspond to negative β coefficients. Statistical significance for all analyses was noted when p-values were less than 0.05, and p-values between 0.05 and 0.06 were noted as trending toward significance.

## Results

### Control group

Our control group was created by combining the animals that were infused with the eGFP virus, but also animals that were originally assigned to the eGFP group but wherein the histological analysis revealed either no-expression (n=9) or unilateral expression (n=4) compared with bilateral expression of eGFP (n=9). To assess if these groups could be collapsed in further analyses, we performed a GLM for each phase of the experiment. Using the mean of the last 3 sessions of training, a GLM was conducted for mean preference using sex and control group (eGFP, unilateral, or no-expression) as between-subject factors, and individual rat as a random factor (full model: *γ ∼ [1 + sex * control group + (1 | rat)*]. We found no significant effect of control group (*p*= 0.54), sex (*p* = .335), or control group * sex interaction (*p*= .955). Similarly, a GLM was performed on *preference change* during the stable preference phase of the experiment, adding drug as a within-subject factor (full model: *γ ∼ [1 + drug * sex*control group + (1 + drug| rat)*]. We found no significant predictors or interactions. Based on these results, the animals in the control groups were collapsed and treated as a single group for subsequent analysis, and added to the ‘virus’ factor as a fourth group (ACC hM4Di, BLA hM4Di, OFC hM4Di, and control).

### Training: Rats developed a consistent preference, responded quickly during choice trials and responded more to the informative cue (S+)

#### Choice

Training data for preference for the info option were first analyzed across all sessions, to evaluate learning of preference on the task. A GLM was conducted for preference for the Info option using sex as between-subject factor, session as within-subject factor, and individual rat as a random factor using the following formula: *γ ∼ [1 + sex*session + (1+session| rat)*]. We found a main effect of session (GLM: β_session_= -0.008, t(868)=-2.09, *p* =0.03) and sex (GLM: βsex= -0.12, t(868)=-2.34, *p* =0.02) but no significant interaction (GLM: β_session_*sex= 0.006, t(868)=1.13, *p* =0.26) (see **Supplement 1**). A main effect of session indicates the preference changed across training, wherein animals begin their preference for information around a probability of 0.5 (indifference) before switching to a preference for the Info or No- Info alternative. While the main effect of sex illustrates that preference between male and female rats differed, the learning rates did not differ by sex given the non-significant interaction of sex by session.

To evaluate wether the observed preference remained consistent within each animal, we compared preference in no-drug conditions before any drug experience. To do so, the last 3 sessions of preference data for training were averaged, and compared to the preference in the last three sessions of the baseline after surgery. A GLM comparing the averaged preference during training and baseline phase was performed, in which phase (training and baseline) was a within-subject factor, and sex was a between- subject factor (full model: *γ ∼ [1 + phase * sex (1 + phase| rat)*]. No significant effects (*p*sex= 0.93, *p*phase= 0.26) or interactions (*p*sex*phase= 0.73) were found, suggesting that preference remained stable across conditions once acquired (See **Figure 3A**).

**Figure 3.**
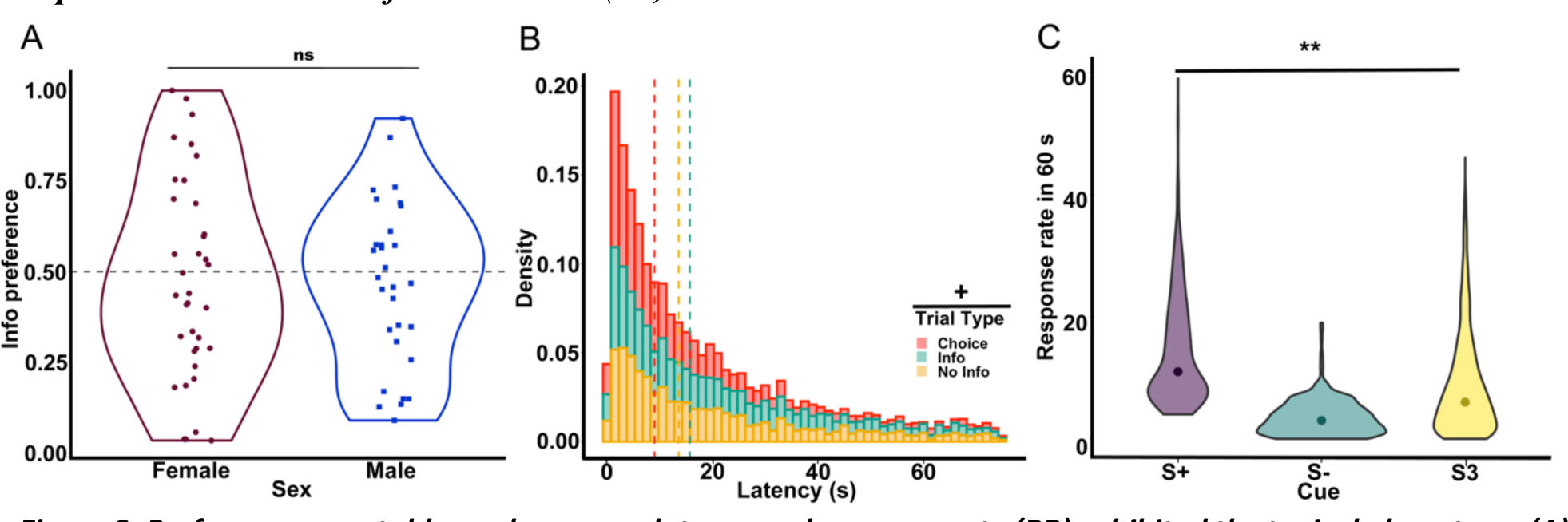
***Preference was stable, and response latency and response rate (RR) exhibited the typical phenotype. (A)*** Mean preference for the info alternative for the last three sessions of training, with individual rats represented as scatter plots. **(B)** Density of response latencies on the last three sessions of training are shown for each trial type. Dashed lines indicate the median latency. A marginally significant difference was found between trial types, where responses in choice trials were faster than in Forced (Info and No Info) trials. **(C)** Violin plots of the distribution of total RR during the 60s cue presentations are presented for each cue during the last three sessions of training. Dots indicate median RR for each cue. A significant effect of cue was found. **p<0.01, +p=0.06, ns = nonsignificant.

#### Latencies

A GLM was conducted on median latencies in the last three sessions of training using trial type (forced info, forced no-info, and choice) as within-subject factors, sex as a between-subject factor, and individual rat as a random factor using the following formula: *γ ∼ [1 + trial type * sex + (1 + trial type| rat)*]. The results yielded a trend for an effect of trial type (GLM: β_trial type_= 6.16, t(179)=1.85, *p* =0.06) wherein rats tended to be faster during choice trials (MedianChoice= 10.6 s, SEM±1.69) than either of the forced trials (MedianForced Info=21.8 s, SEM±3.97, and MedianForced No-Info= 18.1 s, SEM±3.81) (See **Figure 3B**). This result replicates what previous literature has shown in this task in pigeons and starlings (González, Izquierdo et al. 2023).

#### Response Rate

The median response rate (RR) during the 60 s cue was analyzed for the last 3 sessions of training. A GLM was conducted for median RR using sex as between-subject factor, and individual rat and cue (S+, S- and S3) as random factors using the following formula: *γ ∼ [1 + sex + (1|rat:cue)*]. Cue was introduced as a random factor to account for the fact that rats were presented with different frequencies of cues given the programmed contingencies (e.g., S+ present only 20% of all Info trials) but this difference also depended on the rat’s choice. For instance, an animal that chooses the No- Info alternative exclusively during choice trials would have more presentations of cue S3 than any other cue. We found a main effect of cue (GLM: β_cue_= -2.11, t(185)=-2.89, *p* = 0.004), indicating that RR was greater for the cue predicting food (MedianS+= 11 presses/minute, SEM±0.34) than to the no-info cue (MedianS3= 6 presses/minute, SEM±0.14), but the lowest RR was to the cue predicting absence of food (MedianS-= 3 presses/minute, SEM±0.06). This is also in line with previous work in which the lowest RR was reported for S-, and typically a greater RR for S+ than S3 (Hinnenkamp, Shahan et al. 2017, Gonzalez and Blaisdell 2021).

### Info Preference: Inhibition of ACC destabilized preference in female but not male rats

#### Choice

Stable preference was analyzed by averaging the three sessions of baseline, and then comparing this choice preference with the 3 sessions of CNO and three sessions of VEH for each subject. An intial analysis on actual preference was performed (see **Supplement 2**) during Baseline (BL), CNO, and VEH conditions for each group (ACC, BLA, OFC and Control) and sex. The formula of the GLM was: *γ ∼ [1 + group * drug * sex + (1 + drug| rat)*]. We found no effect of any factor (p > 0.41) nor interactions (p > 0.26), probably due to the high variability in initial preference within each group, and therefore, any potential changes in preference depended on each rat’s baseline. Thus, we compared changes in stable preference by calculating the absolute difference between preference in baseline-to- CNO and baseline-to-VEH conditions (See **Figure 4A**). A GLM was conducted for *absolute preference change* using drug as a within-subject factor, virus and sex as between-subject factors, and individual rat as random factor using the following formula: *γ ∼ [1 + virus * drug * sex + (1 + drug| rat)*]. We found a significant interaction of drug*virus (GLM: β_drug*virus_= 0.065, t(124)=4.88, *p* = 3.24^e-06^) and sex*drug*virus (GLM: β_sex*drug*virus_ = -0.065, t(124)=-3.15, *p* = 0.002). Thus, we were justified to conduct follow-up analyses with individual group comparisons: *γ ∼ [1 + virus * drug * sex + (1 + drug| rat)*]. For the comparison of ACC hM4Di vs Control, we found a significant interaction of drug*virus (GLM: β_drug*virus_= 0.074, t(68)=5.58, *p* = 4.61^e-07^) and drug*virus*sex interaction (GLM: β_sex*virus_*drug = -0.077, t(68)= -3.70, *p* = 0.0004). Comparisons of OFC hM4Di vs. Control and BLA hM4Di vs. Control were not significant. Post-hoc analysis using Bonferroni correction and sex as a covariate for the ACC hM4Di group resulted in a significant effect of drug (t(29)= 3.66, *p* =0.002). In contrast, there was no significant effect of drug in the control group (t(41)= 1.59, *p* = 0.239). Overall, these results indicate that ACC inhibition causes a desestabilization of preference in female rats.

**Figure 4.**
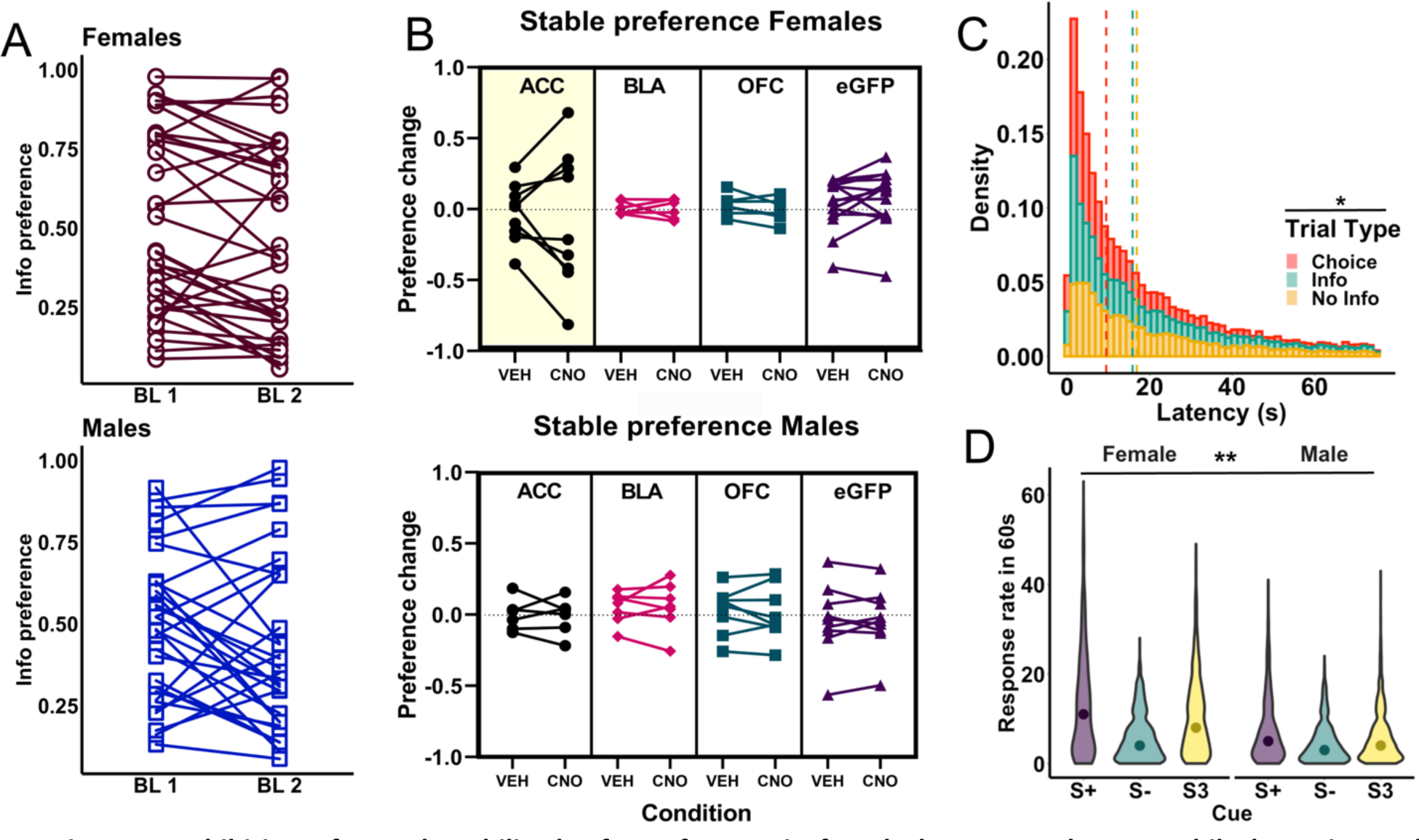
Inhibition of ACC destabilized Info preference in female but not male rats, while latencies and response rate (RR) were unaltered by inhibition. (A) Baseline Info preference for females (top) and males (bottom) over two baselines. The first baseline (BL1) was obtained after surgery and the second baseline (BL2) was obtained at the end of the experiment. Note the lack of dramatic changes within individual animals. **(B)** Mean change in preference for the info alternative between the last three sessions of baseline after surgery minus the preference during VEH and CNO administration for females (top) and males (bottom). Note that a positive change indicates a shift in preference towards the non-info option, whereas a negative change indicates a preference shift toward the info option. Values around 0 indicate no change in preference. **(C)** Density of latencies to choose across drug conditions is shown for each trial type. Dashed lines indicate the median latency. A significant difference between trial types was found, where choice trials were faster than Forced (Info and No Info) trials. **(D)** Violin plots of the distribution of total RR during the 60s cue presentations are presented for each cue for all drug conditions for females (left) and males (right). Dots indicate median response rate for each cue. Females responded more than males and a significant effect of cue was found. **p<0.01, *p<0.05.

#### Latencies

Latency to choose was analyzed by calculating the median for CNO and VEH sessions (See **Figure 4B**). A GLM was conducted for median latency using drug and trial type as within-subject factor, virus and sex as between-subject factors, and individual rat as a random factor using the following formula: *γ ∼ [1 + trial type * virus * drug * sex + (1 + trial type + drug| rat)*]. We found a main effect of trial type (GLM: β_trial type_= 5.61, t(380)=2.19, *p* =0.028) indicating that rats were faster responding during choice trials (MedianChoice= 8.4 s, SEM±0.4) than forced trials (MedianForced Info=14.8 s, SEM±0.66, and MedianForced No-Info= 15.1 s, SEM±0.62). Note that we did not find any difference by sex or virus, suggesting that the change in preference observed in the female rats following ACC inhibition did not affect decision speed.

#### Response rate

RR during the 60s cue duration was also analyzed by calculating the median during CNO and VEH sessions (See **Figure 4C**). A GLM was conducted for median RR using drug as within-subject factor, virus, and sex as between-subject factors, and individual rat and cue (S+, S- and S3) as random factors using the following formula for the full model: *γ ∼ [1 + cue * virus * drug * sex + (1 + drug| rat:cue)*]. As in training, cue was defined as a random factor given that there were different probabilities in experiencing each cue based on an individual rat’s choices. We found a main effect of sex (GLM: β_sex_ = -7.87, t(379)=-2.17, *p* = 0.03), with females showing higher response rates (MedianFemale= 7 presses/min, SEM±0.06) than males (MedianMale= 4 presses/min, SEM±0.46).

### Previous choices are not as predictive of current information choices in ACC-inhibited females

To further understand the nature of the effect of ACC inhibition on choice behavior, we fit a logistic regression, a type of GLM for binary classification, to each rat’s data that belonged to one of the experimental groups (ACC hM4Di-CNO, BLA hM4Di-CNO, OFC hM4Di-CNO), or control group-CNO for females and males. Using the three features described in the analysis section we aimed to predict the probability of choice. We computed the odds ratio and 95%, 80% CI for each condition’s regression fit: If an odds ratio was greater than 1, it represented an increase in the odds of the outcome happening given a one-unit increase in the predictor. If the odds ratio was less than 1, it represented a decrease in the odds of the outcome happening given a one-unit increase in the predictor. And finally, an odds ratio of exactly 1 signified that the value of the predictor did not affect the probability of the outcome.

The model resulted in a significant predictor of Previous Choice to current choice for females and males across all groups (*tACC-fem*(392)= 5.19, *p* = 2.05^e-07^; *tBLA-fem*(2385)= 6.55, *p* = 5.66^e-11^; *tOFC-fem*(249)= 7.05, *p* = 1.77^e-12^; *tcontrol-fem*(331)= 23.64, *p* = 1.23^e-123^; *tACC-male*(314)= 8.48, *p* = 2.22^e-17^; *tBLA-male*(1277)= 8.26, *p* = 1.39^e-16^; *tOFC-male*(266)= 8.31, *p* = 8.80^e-17^; *tcontrol-male*(334)= 17.12, *p* = 9.02^e-66^), however, it is important to note that the ACC female group exhibited a lower odds ratio, indicating that when ACC was inhibited the previous choice became a weaker predictor of current choice than in the other groups.

Previous Reward | Info was also a significant predictor of current choice for females and males across groups (*tACC-fem*(392)= -2.28, *p* = 0.02; *tBLA-fem*(2385)= -5.61, *p* = 1.92^e-08^; *tOFC-fem*(249)= -6.05, *p* = 1.42^e-09^; *tcontrol-fem*(331)= -17.85, *p* = 2.49^e-71^; *tACC-male*(314)= -6.11, *p* = 9.86^e-10^; *tBLA-male*(1277)= -5.08, *p* = 3.68^e-07^; *tOFC-male*(266)= -5.56, *p* = 2.63^e-08^; *tcontrol-male*(334)= -13.59, *p* = 4.35^e-42^), but Previous Reward|No Info and Latency were not significant predictors of current choice (*p* > 0.19) (**Figure 5**). Given that we found that the effect of previous reward depended on which option was chosen in the trial, we calculated the percentage of switching if rats received a reward for the Info option (that is, switching to the No-Info, given that they selected the Info option and received a reward): 72.15% for females and 86.95% for males across all groups. This pattern was not observed when rats received a reward after choosing the No-Info option (17.85% for females and 49.84% for males across conditions). This result indicates adoption of a *win-stay* strategy following NoInfo choices but a *win-switch* strategy following Info choice for ACC-inhibited females. Note that this is different when we ignore reward, for choice alone (PrevChoice, **Figure 5**): the percentage of switch in Males is 18.38% and 27.38% for Info and No-Info choices, respectively, and 25.47% and 27.42% in Females for Info and No-Info choices, respectively.

**Figure 5.**
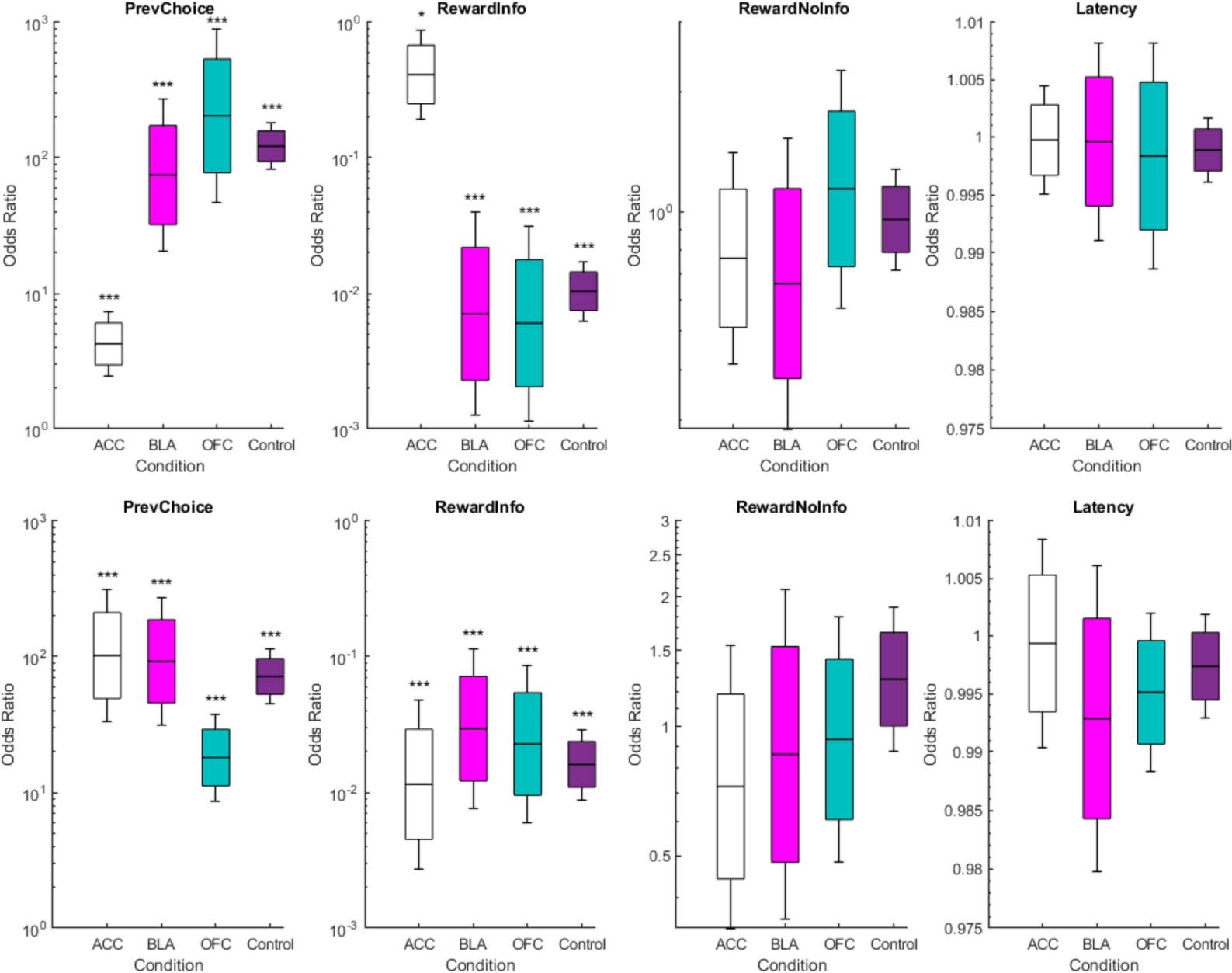
***Female, but not male, rats adopt a win-switch strategy following rewarded Info trials during ACC inhibition.*** Each panel represents a feature of the model (Previous Choice, Previous Reward in an Info choice, Previous Reward in a No-Info Choice, and Latency) for female (top) and male (bottom) animals following CNO administration in four different groups: ACC hM4Di, BLA hM4Di, OFC hM4Di and Control. The y-axis is the odds ratio of the predictor. Boxes, error bars, and lines represent 80% confidence interval, 95% confidence interval, and mean odds ratio, respectively. For females: Mean model R^2^ for control group = 0.4613, ACC group = 0.4189, OFC group = 0.4613, BLA group = 0.5181. For males: Mean model R^2^ for control group = 0.0912, ACC group= 0.4280, OFC group = 0.5275, BLA group= 0.3971. *p < 0.05, **p < 0.001, ***p < 0.0001.

These findings suggest that previous choice has the most infuence on subsequent behavior for all conditions, however, we found a low odds ratio for the ACC females but not ACC males, indicating that previous choice was not as reliable a predictor for the former. These results also shed light on the mechanism behind the reported effect of ACC inhibition in females, suggesting that female rats used a paradoxical win-switch strategy following rewarded Info trials. We further discuss the interpretation of these results below.

## Discussion

Despite the well-characterized roles of OFC and ACC in decision-making (Izquierdo 2017, Bromberg-Martin and Monosov 2020, Sosa, Buonomano et al. 2021), few studies find clear dissociations between these regions in rodents. For example, both OFC and ACC are important in confidence report as measured by temporal wagering (Stolyarova et al. 2019; Lak et al. 2014) and are also involved in stimulus-based reversal learning (Ye et al. 2023). In the present study, we found a clear dissociation between ACC and OFC that may shed light on their individual contributions in decision-making about non-instrumental information, and point to a hierarchy of functions within rodent frontal cortex.

Specifically, we found that ACC inhibition rendered animals’ decisions about information more stochastic, also corroborated by the logistic regression model which showed that previous choice was not a good predictor of future choices when ACC was offline. The pattern of results on latencies and response rate also indicates the effect of inhibition on decision-making is not due to performance decrements (i.e., we found unchanged latencies and response rates). Interestingly, the logistic regression revealed a similar finding of reduced use of previous choice following OFC inhibition, but this did not translate into a significant change in behavior. Thus, it could be that “performance monitoring” is a general feature of rodent frontal cortex, with ACC exerting a more prominent role than OFC. Indeed, ACC has been linked more to the representation of reward opportunities across the environment at different timescales while also keeping track of animals’ previous actions (Kolling, Behrens et al. 2016, Wittmann, Kolling et al. 2016, Spitmaan, Seo et al. 2020), suggestive of a special metacognitive role (van Veen, Holroyd et al. 2004, Stolyarova, Rakhshan et al. 2019, Kane, James et al. 2022, Takeuchi, Roy et al. 2022).

Our results presented here also replicate the vast literature investigating the behavior behind the preference for the informative, yet “suboptimal” alternative. As other researchers have previously reported, the preference for the informative option is more variable in rats than pigeons or starlings when that option results in less food (Stagner and Zentall 2010, Vasconcelos, Monteiro et al. 2015). This indicates that perhaps there are different sensitivities to information and/or reinforcement history between species. Regarding latencies to make a choice, we replicate what has been observed in several studies: animals respond faster on “true” choice trials than forced-choice trials. Ecologists have discussed this counterintuitive result by suggesting that animals did not evolve to make simultaneous but rather sequential choices in nature, such that latencies during forced choice trials can be used to predict preference (Kacelnik, Vasconcelos et al. 2010). Finally, response rate during the cue presentation followed the typical pattern reported previously: animals respond more in the presence of the always and partially-reinforced cue, and do not respond to the non-reinforced cue (Hinnenkamp, Shahan et al. 2017, Gonzalez and Blaisdell 2021). It is important to highlight that animals are not required to respond during the duration of the cue; however, they typically do, suggesting a Pavlovian association established between the cue and the outcome. This association we believe is central to the preference, in which a stable preference emerges when an association between the reinforced cue (S+) and the outcome and the non-reinforced cue (S-) and the outcome is established (González, Izquierdo et al. 2023).

A similar variation of the task presented in this study was used to assess preference for non-instrumental information in humans and monkeys. In those studies, subjects are given a choice between two alternatives: one that provides informative cues that indicate the trial’s outcome, and another provides non-informative cues that do not indicate the trial’s outcome. Similar to our task, the information provided by the cues does not influence or change the outcome. However, humans and monkeys were also informed about the quantity (money for humans and juice for monkeys) of the outcome in a given trial. The results showed that macaque monkeys and humans prefer information, and that they are willing to sacrifice water/money to obtain immediate information about the outcomes. One group previously found that OFC neurons encode variables that are relevant in learning and decision-making but do not integrate these variables into a single value (Blanchard, Hayden et al. 2015, Aguirre, Woo et al. 2023).

This result is consistent with our findings in which OFC inactivation resulted in an attenuated use of previous choice information revealed by the logistic regression, suggesting problems in retrieving the memory of previous choice, but this did not translate into a real change in preference. ACC may have a higher-level role in sustaining information-seeking (Hunt, Malalasekera et al. 2018, White, Bromberg-Martin et al. 2019). For example, activity ramps up in this region in anticipation of information becoming available to resolve uncertainty before reward delivery (White, Bromberg-Martin et al. 2019). Our results support the monitoring role of ACC in decisions about information, where inhibition altered the integration of value, destabilizing preference.

We found that ACC inhibition desestabilized preference, but only in female animals. We did not find changes in latency to choose or response rates, indicating that the observed differences were not due to motor impairment or due to changes in the association between cues and outcomes. Therefore, the change in preference with ACC inhibition is unlikely due to problems in accessing overall value of each option. Previous studies have determined that ACC does not support simple effort or the hedonic value of a given alternative (i.e., one option). Instead, it computes the value *across* options, that is, it tracks the overall “better” choice (Hart, Gerson et al. 2017, Hart, Blair et al. 2020). Here, we found that the change in preference following ACC inhibition was not in any particular direction. If animals with ACC offline had issues in accessing the overall value of the best option, in this case the no-info option, we would have observed an increase in preference for the info option only. In contrast, we found that preference became more stochastic. Previous research has shown that ACC needs to be engaged when animals are asked to stick to a strategy (i.e., win-stay) and animals instead show high variability of responses, thus more stochastic behavior, when ACC is offline (Tervo, Proskurin et al. 2014, Tervo, Kuleshova et al. 2021).

The preference change may also indicate a reduced ability to link previous actions (in this case, the previous lever press) to the current trial. However, this is unlikely because the motor response to the lever press did not change (i.e., latencies were unaffected), nor did the assigned value of the cues (i.e., response rate to cues also did not change). We propose instead that ACC inhibition results in an impairment in accessing the value of information. Previous studies using this paradigm indicates that the contrast between both informative cues (i.e., the difference in information between the S+ and the S-) is essential for the development of a preference for information. Other research has already suggested that ACC is important for information-seeking behavior; however, these studies reported this following mosly electrophysiological recording and neuroimaging in monkeys and humans, respectively (Kennerley and Wallis 2009, Monosov 2017, Bromberg-Martin and Monosov 2020). Our study provides the first causal evidence of the importance of ACC in decision-making involving information.

Sex differences in the involvement of ACC in value-based decision making have been reported before. In a recent study, Cox et al. (2023) found that ACC inhibition disrupted the relationship between the value of each alternative and motivation to engage in the task in female mice. However, they did not find changes in preference as we found in our study. This group also reported that the ACC-to-dorsomedial striatal neuron pathway represents negative outcomes more strongly in female than males rats, suggesting differential sensitivity to negative feedback (Cox, Minerva et al. 2023). Similarly, in tasks involving decision-making under risk, it has been determined that female rats are more risk-averse (Orsini, Willis et al. 2016), indicating that female and male rats use different strategies to make decisions (Orsini, Blaes et al. 2021). In our task, similar to Cox et al. (2023), we did not find differences in preference between female and male rats before inhibition. However, the destabilization of preference might indicate that females rely more strongly on ACC to keep track of reward statistics.

Similar to the results in rodents, groups studying human subjects have uncovered sex differences in strategy in decision-making (Chowdhury, Wallin-Miller et al. 2019, Chen, Knep et al. 2021), usually in tasks involving risk (van den Bos, Homberg et al. 2013). These sex differences are important to understand because, on the one hand, there is a higher prevalence of depression or anxiety-related disorders in women (Cyranowski, Frank et al. 2000); and on the other hand, there is increasing evidence that neuropsychiatric conditions such as ADHD, bipolar disorder, and autism show different onset, symptom severity, and prognosis dependent on sex (Grissom and Reyes 2019, Hwang, Arnold et al. 2020). This evidence suggests differential involvement of circuits involved in decision-making by sex, and our study contributes to the effort in finding the neural mechanisms behind these potential differences.

## Acknowledgment

We thank Dr. Pamela Mattar for her help with the protocol and technique for c-fos IHC, and members of the Izquierdo lab for early feedback and suggestions on these experiments.

## Author contributions

**ABP:** Resources, funding, writing- review & editing. **AI:** Conceptualization, methodology, formal analysis, data curation, resources, writing-original draft, writing- review & editing, software, supervision, and funding. **AR:** Investigation and data curation. **IY:** Investigation and data curation. **JLR:** Investigation, writing- review & editing. **SA:** Investigation, writing- original draft, and data curation. **VVG**: Conceptualization, methodology, formal analysis, investigation, data curation, writing- original draft preparation, writing- review & editing, software, visualization, and funding. **YZ:** Software and formal analysis.

