## Supplementary Figures for "A special role for anterior cingulate cortex, but not orbitofrontal cortex or basolateral amygdala, in choices involving information"

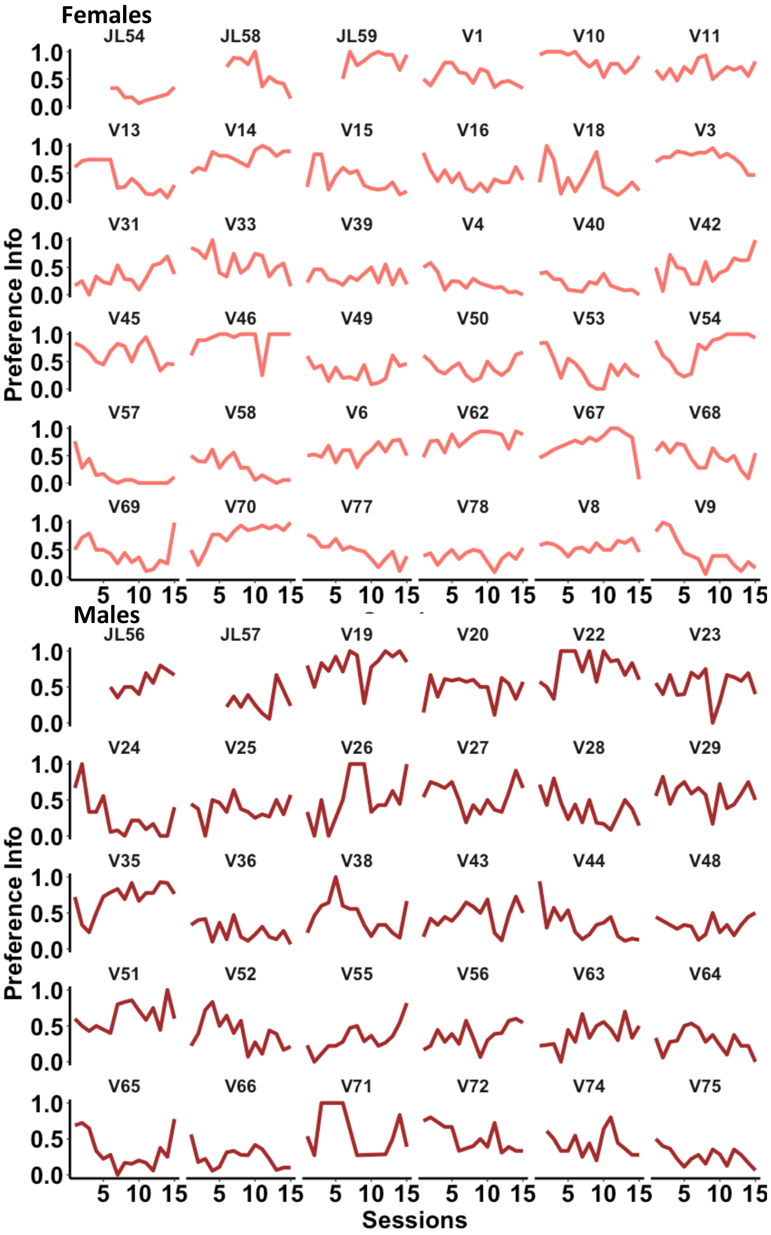


***Supplement 1. Acquisition curves during initial training by rat.*** Rats received 15 sessions of training prior to the stereotaxic surgery to facilitate learning and testing. Color indicates the sex of the animal.

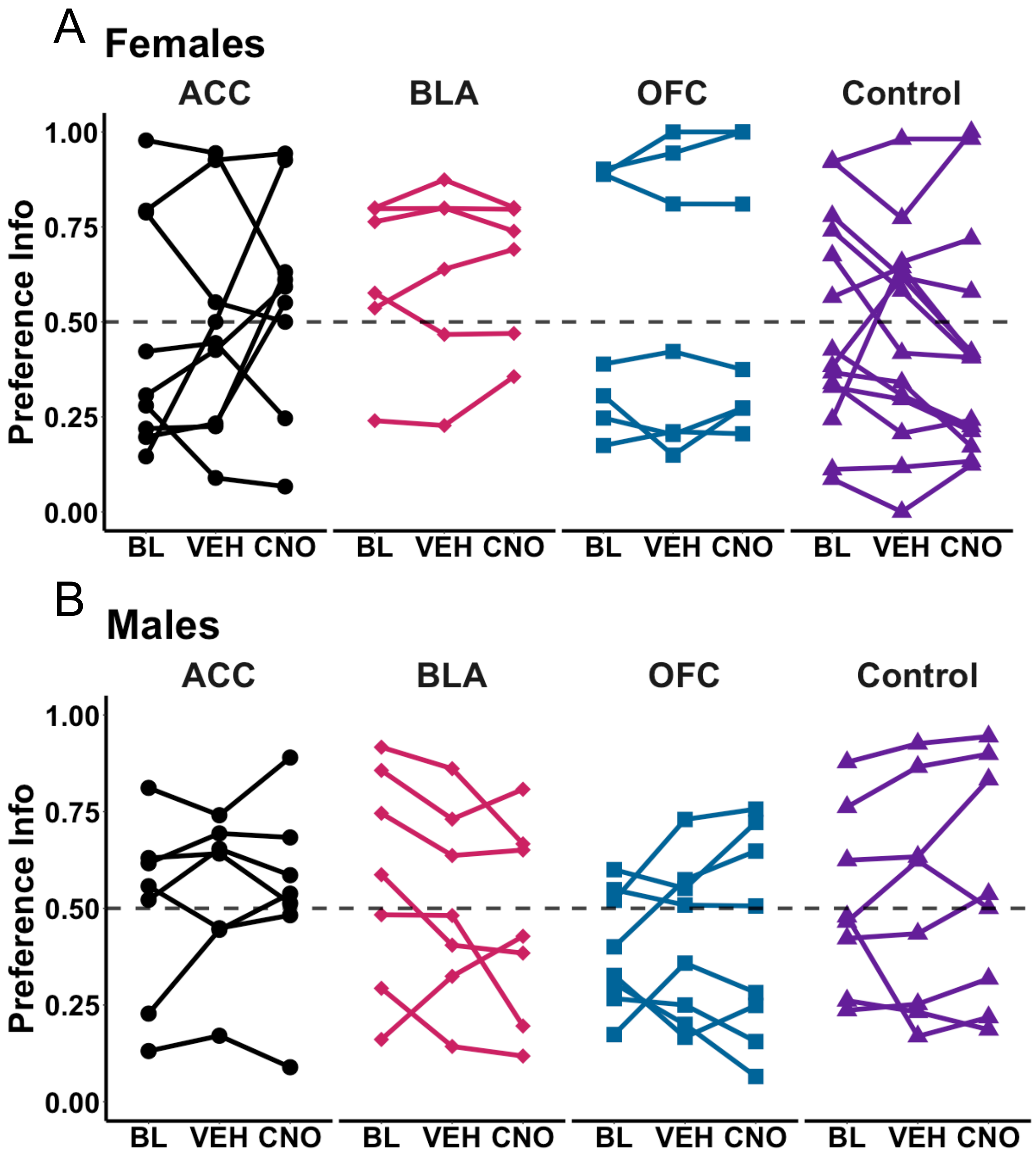


***Supplement 2. Preference during stable performance by group and drug condition.*** The average during No drug (Baseline, BL), vehicle (VEH) and CNO sessions are presented by Group (ACC, BLA, OFC and Control) for female (**A**) and male (**B**) rats.

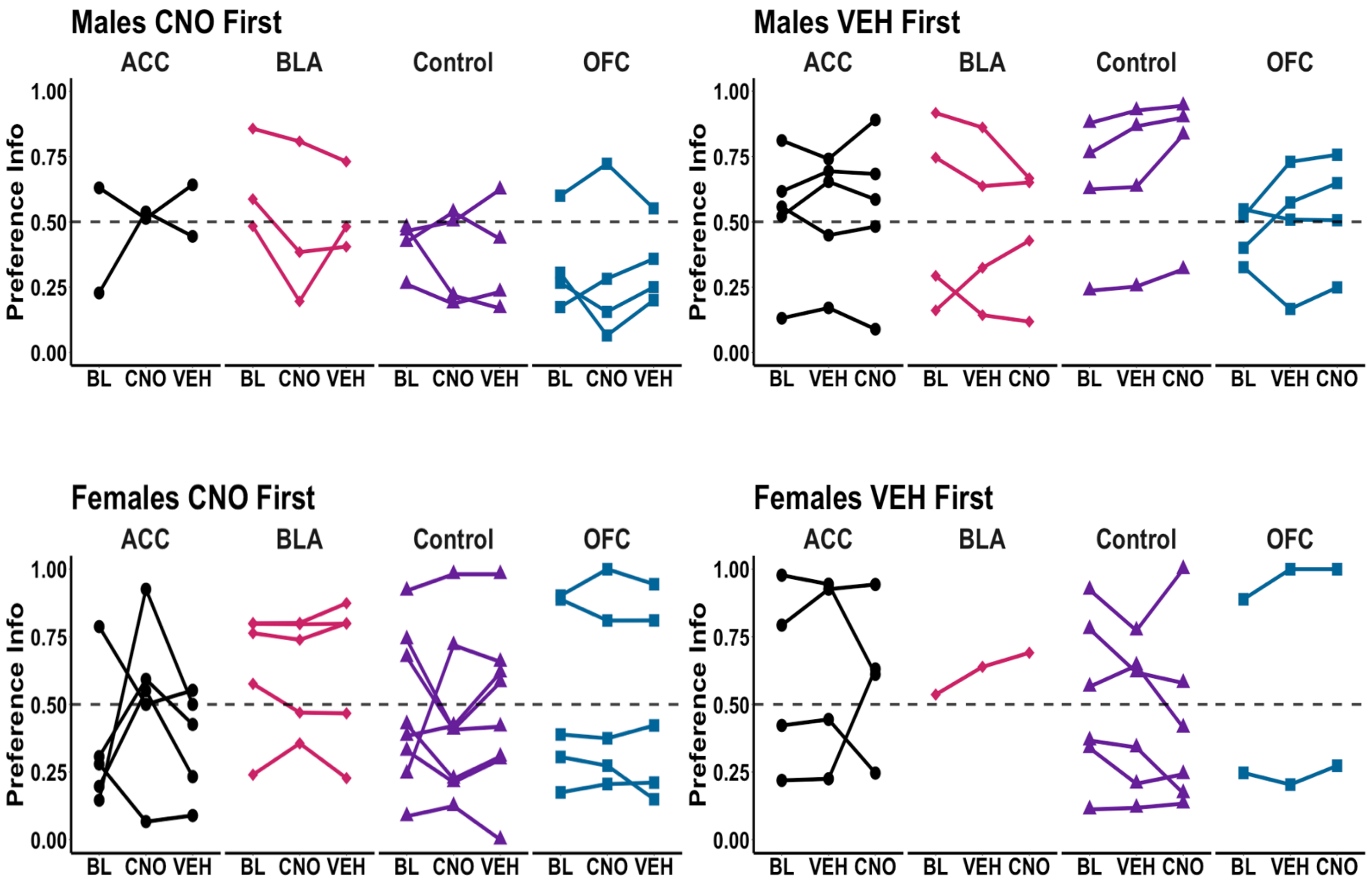


***Supplement 3. Preference during stable performance separated by order of drug administration.*** The average during No drug (Baseline, BL), vehicle (VEH) and CNO sessions are presented by Group (ACC, BLA, OFC and Control) for male (top) and female (bottom) rats. The conditions in the x axis indicate the order in which each rat receive the conditions.
